# Functional role of lanthanides in enzymatic activity and transcriptional regulation of PQQ-dependent alcohol dehydrogenases in *Pseudomonas putida* KT2440

**DOI:** 10.1101/140046

**Authors:** Matthias Wehrmann, Patrick Billard, Audrey Martin Meriadec, Asfaw Zegeye, Janosch Klebensberger

## Abstract

The oxidation of alcohols and aldehydes is crucial for detoxification and efficient catabolism of various volatile organic compounds (VOCs). Thus, many Gram-negative bacteria have evolved periplasmic oxidation systems, based on pyrroloquinoline quinone-dependent alcohol dehydrogenases (PQQ-ADHs), which are often functionally redundant. Using purified enzymes from the soil-dwelling model organism *Pseudomonas putida* KT2440, the present study reports the first description and characterization of a lanthanide-dependent PQQ-ADH (PedH) in a non-methylotrophic bacterium. PedH exhibits enzyme activity on a similar substrate range as its Ca^2+^-dependent counterpart PedE, including linear and aromatic primary and secondary alcohols as well as aldehydes, however, only in the presence of lanthanide ions including La^3+^, Ce^3+^, Pr^3+^, Sm^3+^ or Nd^3+^. Reporter assays revealed that PedH not only has a catalytic function, but is also involved in the transcriptional regulation of *pedE* and *pedH*, most likely acting as a sensory module. Notably, the underlying regulatory network is responsive to as little as 1 – 10 nM of lanthanum, a concentration assumed to be of ecological relevance. The present study further demonstrates that the PQQ-dependent oxidation system is crucial for efficient growth with a variety of volatile alcohols. From these results we conclude that functional redundancy and inverse regulation of PedE and PedH represents an adaptive strategy of *P. putida* KT2440 to optimize growth with volatile alcohols in response to different lanthanide availability.

**IMPORTANCE:** Due to their low bioavailability, lanthanides have long been considered as biologically inert. In recent years however, the identification of lanthanides as a cofactor in methylotrophic bacteria has attracted tremendous interest among various biological fields. The present study reveals that one of the two PQQ-ADHs produced by the model organism *P. putida* KT2440 also utilizes lanthanides as a cofactor, thus expanding the scope of lanthanide employing bacteria beyond the methylotrophs. Similar to methyloptrophic bacteria, a complex regulatory network is involved in the lanthanide-responsive switch between the two PQQ-ADHs encoded by *P. putida* KT2440. We further show that functional production of at least one of the enzymes is crucial for efficient growth with several volatile alcohols. Overall, our study provides a novel understanding for the redundancy of PQQ-ADHs observed in many organisms and further highlights the importance of lanthanides for bacterial metabolism, particularly in soil environments.

## INTRODUCTION

As a soil-dwelling organism, *Pseudomonas putida* can encounter a large diversity of volatile organic compounds (VOCs) from different sources (1–3). The ecological role of many VOCs is not clearly defined but the number of known specific functions is rapidly increasing. These functions include the growth promotion of plants, anti-herbivore, -bacterial, and -fungal activities, and signaling both within the same and between different species (4–7). Among many other chemicals, VOCs include cyclic, acyclic, aromatic, and terpenoid structures with alcohol and aldehyde moieties, which are mainly derived from the metabolism of bacterial, yeast, fungal, or plant species. Beside their specific molecular function, they can also serve as carbon and energy sources for a wide range of microorganisms. To efficiently use volatile alcohols and aldehydes, it is advantageous if their metabolism is initiated by pyrroloquinoline quinone (PQQ)-dependent alcohol dehydrogenases (PQQ-ADHs) for at least two different reasons. Firstly, by using a periplasmic oxidation system, the organism is able to rapidly detoxify the often harmful chemicals without prior transport into the cytoplasm (8, 9). Secondly, the periplasmic location of the enzymes allows the rapid capture of a volatile carbon source by the conversion to and accumulation of acidic products with decreased volatility.

The reaction mechanism of PQQ-ADHs is still not completely resolved, but most likely proceeds *via* an ion-assisted direct hydride transfer from the substrate to the C5 of the non-covalently bound cofactor PQQ (10–12). PQQ-ADHs can be divided into different subclasses depending either on their molecular composition (quinoproteins or quinohemoproteins), or whether they are membrane bound or freely soluble within the periplasm (13). Many organisms express different classes or even multiple PQQ-ADHs of the same type, indicating the importance of these enzymes (14–16). The genome of *P. putida* KT2440 encodes two PQQ-ADHs, namely PP_2674 *(pedE)* and PP_2679 *(pedH)*, which have been shown to be involved in the metabolism of different substrates (17–19). PedE is a homolog of ExaA (QEDH) from *Pseudomonas aeruginosa*, which represents the most intensively studied member of the class of soluble ethanol dehydrogenases (20–24). ExaA and homologs thereof accept a wide variety of substrates and rely on a Ca^2+^ ion in the active site, in addition to the PQQ cofactor, for the oxidation of primary and secondary alcohols, as well as aldehydes (18, 24). Despite their broad substrate range, ExaA-like enzymes show only very poor conversion of methanol. Not surprisingly, methano- and methylotrophic bacteria, which can use methane and methanol as a source of carbon and energy, encode a different type of PQQ-dependent enzyme, the MxaF-type of methanol dehydrogenases (MxaF-MDH) (25, 26). These enzymes display high substrate specificity for methanol and formaldehyde and also depend on a Ca^2+^ as cofactor (27). Interestingly, methano- and methylotrophic bacteria encode an additional type of PQQ-dependent methanol dehydrogenases, the XoxF-MDH type, which utilizes rare earth metals (REM) of the lanthanide series as cofactor instead of calcium (28–30).

Since their discovery, several XoxF-type MDHs from different methano- and methylotrophs have been identified and characterized (31–33). Phylogenetic analysis of available sequence information suggests that lanthanide dependency is an ancestral feature of PQQ-ADHs, and that these enzymes are more abundant than their Ca^2+^-dependent counterparts (15, 34). In addition, a very recent publication described the first lanthanide-dependent ethanol dehydrogenase in *M. extorquens* AM1 (16). As a consequence, REM-dependent enzymes and the microorganisms which produce them have sparked a lot of academic and commercial interest, as they might be exploited in a broad variety of biotechnological fields (35, 36). Potential applications range from the development of new biocatalysts and biosensors to the use of the associated microorganisms in biomining, bioleaching, and recycling processes for REMs. However, so far lanthanide-dependent PQQ-ADHs have been limited to methano- and methylotrophic bacteria.

In the present study, we report the first description and detailed characterization of a lanthanide-dependent PQQ-ADH (PedH) in the non-methylotrophic bacterium *Pseudomonas putida* KT2440, which represents a model organism for industrial and environmental applications (37–40). We demonstrate that PedH exhibits enzymatic activity only in the presence of lanthanides, including but not limited to lanthanum, praseodymium and cerium, and show that this enzyme has a similar substrate range as PedE, the recently characterized Ca^2+^-dependent PQQ-ADH from KT2440 (18). By the use of deletion mutants and transcriptional reporter fusions, we provide evidence that the functional redundancy of the PQQ-ADHs reflects the variable availability of lanthanides in the natural environment of *P. putida* KT2440 and show that these enzymes are crucial for efficient growth with a variety of volatile alcohols. Finally, we reveal that PedH plays an important role in the regulatory switch between *pedH* and *pedE* transcription, most likely acting as a sensory module. From these data, we conclude that KT2440 responds to lanthanide availability with the inverse transcriptional regulation of the two PQQ-ADHs to optimize growth with volatile alcoholic and aldehyde substrates.

## RESULTS

### Biochemical characterization of PedE and PedH

Like many other organisms, *Pseudomonas putida* KT2440 harbors more than one gene annotated as a PQQ-ADH, namely PP_2674 (PedE/QedH; GI: 26989393) and PP_2679 (PedH; GI: 26989398). To study the rationale for this redundancy, we purified and characterized the corresponding enzymes. A one-step affinity chromatography method produced soluble C-terminally His-tagged PedE and PedH to visible purity (**Fig. S1**) from cell lysates of *E. coli* BL21 (DE3). Under optimized reaction conditions, which include the presence of 1 mM Ca^2+^, the specific activities of purified PedE with a variety of substrates were determined (**Table 1**). For all linear primary alcohols and aldehydes, comparably high enzyme activities ranging from 1.9 ± 0.2 U mg^−1^ to 6.7 ± 0.9 U mg^−1^ were found. Similarly, 2-phenylethanol, the secondary alcohol 2-butanol, cinnamyl alcohol, and the acyclic sesquiterpene farnesol were efficiently converted with specific activities ranging from 6.7 ± 1.1 U mg^−1^ to 2.0 ± 0.3 U mg^−1^. Methanol, 2,3-butanediol, and ethanolamine were poor substrates for the enzyme with about 10-fold decrease in specific activity compared to ethanol or 2-phenylethanol. From all substrates tested, cinnamyl aldehyde was the only compound with which no activity was detected for PedE.

**Table 1:** Specific activities of PedE and PedH with various alcohols and aldehydes. Data are presented as the mean value of three independent measurements with the corresponding standard deviation. 10 mM of substrate was used in H_2_O if not indicated otherwise.

| Substrate | Specific activity [U mg<sup>-1</sup>] |  |
| --- | --- | --- |
|  | PedE | PedH |
| Methanol | 0.61 ± 0.10 | 0.80 ± 0.05 |
| Ethanol | 6.7 ± 0.9 | 11.0 ± 0.3 |
| Ethanolamine | 0.55 ± 0.09 | 1.6 ± 0.2 |
| 1-Butanol | 5.8 ± 0.1 | 11.5 ± 0.7 |
| 2-Butanol | 4.4 ± 0.7 | 7.6 ± 0.4 |
| 2,3-Butanediol | 0.39 ± 0.03 | 0.78 ± 0.04 |
| 1-Hexanol | 5.2 ± 0.1 | 10.4 ± 1.1 |
| 1-Octanol<sup>a</sup> | 3.5 ± 0.2 | 4.7 ± 0.7 |
| 2-Phenylethanol | 6.7 ± 1.1 | 10.2 ± 1.4 |
| Acetaldehyde | 4.7 ± 0.5 | 6.7 ± 0.4 |
| Butyraldehyde | 6.1 ± 0.4 | 10.3 ± 0.6 |
| Hexanal<sup>a</sup> | 3.8 ± 0.1 | 6.2 ± 0.3 |
| Octanal<sup>a</sup> | 1.9 ± 0.2 | 3.6 ± 0.6 |
| Cinnamyl alcohol<sup>a</sup> | 2.4 ± 0.1 | 3.9 ± 0.1 |
| Cinnamaldehyde<sup>b</sup> | n. d.<sup>c</sup> | n. d.<sup>c</sup> |
| Farnesol<sup>b</sup> | 2.0 ± 0.3 | 3.8 ± 0.5 |

<sup>a</sup> 10 mM substrate in DMSO.
<sup>b</sup> 500 μM substrate in DMSO.
<sup>c</sup> Activity below detection limit.

When we assayed purified PedH under the optimized reaction conditions used for PedE, no activity for any of the tested substrates was observed (data not shown). Comparison of the active sites of both enzymes, using homology models based on the crystal structure of the ethanol dehydrogenase ExaA of *P. aeruginosa* (PDB: 1FLG) revealed that, similar to other characterized representatives of the PQQ-dependent ethanol dehydrogenase type, the PedE protein harbors a serine residue at amino acid position 295, which is involved in the coordination of the Ca^2+^ ion (**Fig. 1A**). In contrast, in PedH this residue is exchanged to an aspartate (**Fig. 1B**). As this aspartate residue has recently been associated with the coordination of trivalent lanthanide ions in the active site of PQQ-dependent methanol and ethanol dehydrogenases in methylotrophs (15, 16), we tested PedH for activity with ethanol in the presence of a variety of rare earth metals (**Fig. 2**). From these experiments we found that PedH showed no activity when 1 μM of Er^3+^, Sc^3+^, Y^3+^ or Yb^3+^ was added to the reaction mixture. However, in the presence of 1 μM of the lanthanides La^3+^, Ce^3+^, Pr^3+^, Nd^3+^, Sm^3+^, Gd^3+^ or Tb^3+^ enzymatic activities were detected, with maximal specific activities observed with Pr^3+^ and Nd^3+^, and only very low activities with Gd^3+^ or Tb^3+^.

**FIG 1:**
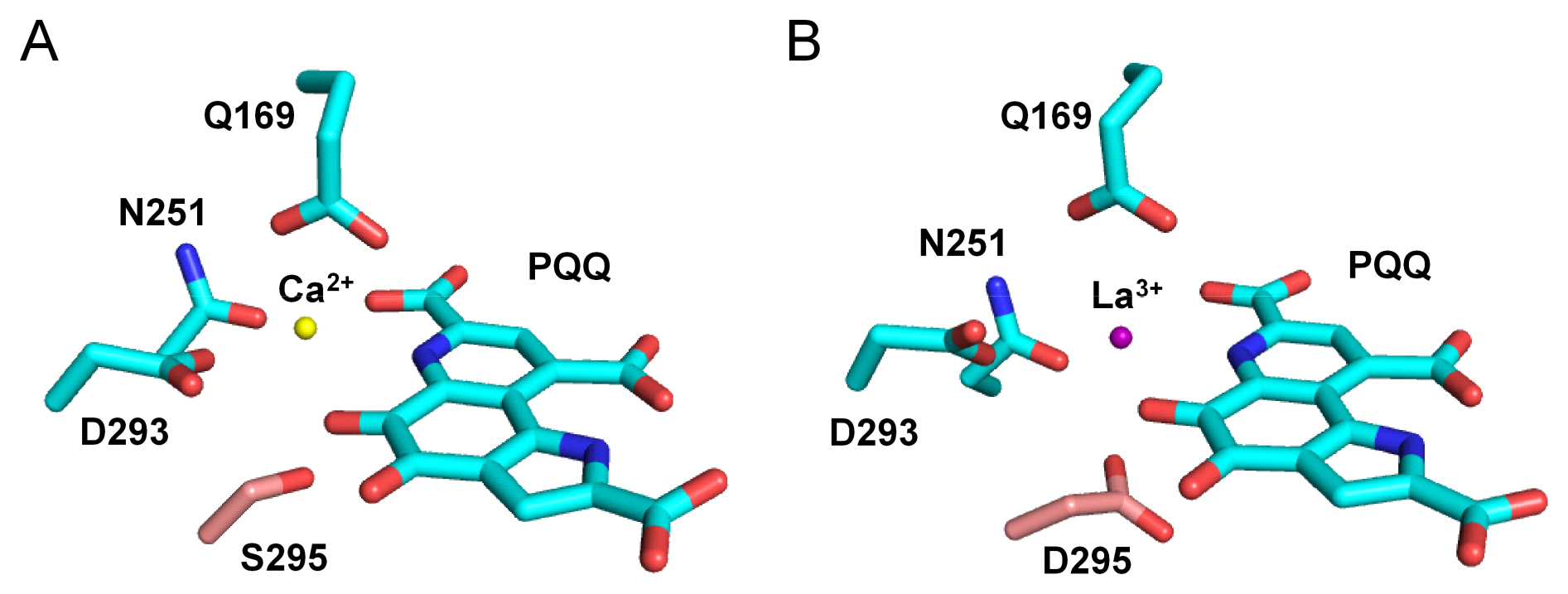
Homology models of PedE (**A**) (GI: 26989393) and PedH (**B**) (GI: 26989398) generated with SWISS-MODEL based on the crystal structure of ExaA from *Pseudomonas aeruginosa* (PDB: 1FLG) using Pymol (70). The catalytic cation *(yellow* or *violet sphere)* coordinating amino acids and the PQQ cofactor are shown as sticks using an element color code (C = cyan, O = red, N = blue). The amino acid position 295 in PedE and PedH is highlighted by using a different color code (C = light red).

**Fig 2:**
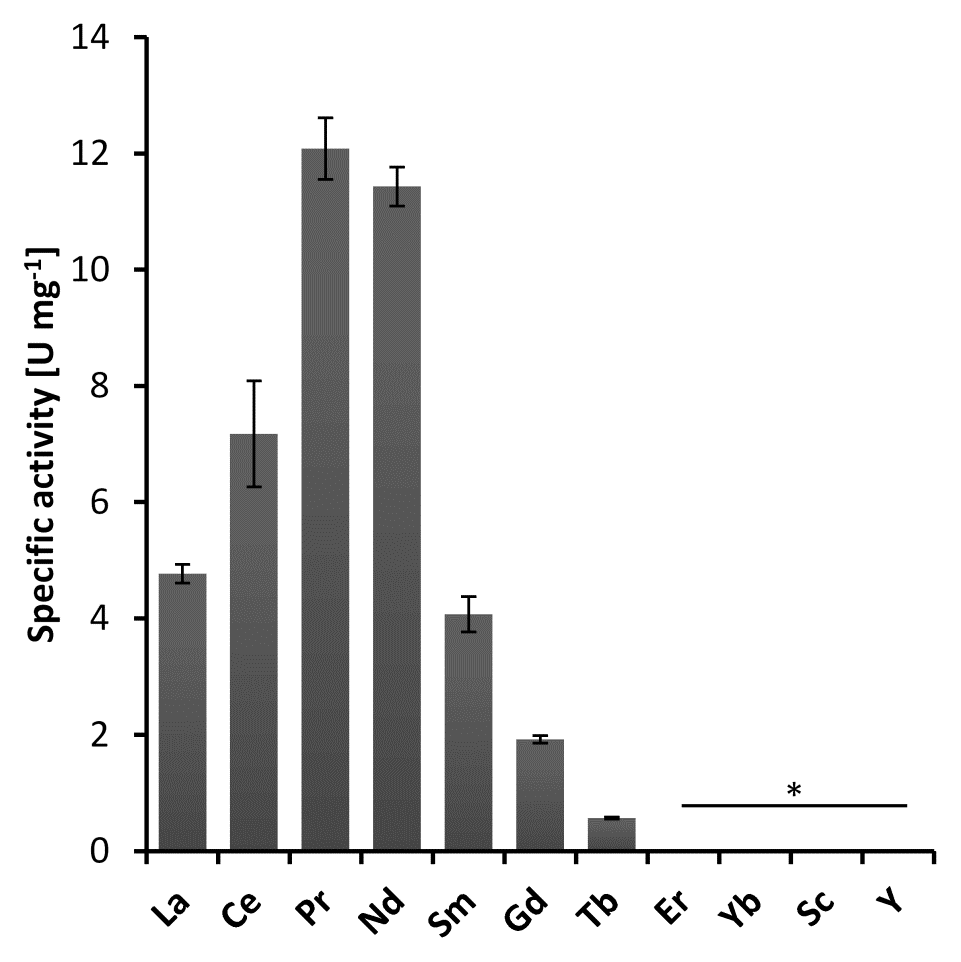
Specific activities of PedH in the presence of 1 μM of various rare earth metal ions with 10 mM ethanol as substrate. Activities below detection limit are indicated (*). Data are presented as the mean value of three replicates and error bars represent the corresponding standard deviation.

Under optimized conditions, which include the supplementation with 1 μM Pr^3+^ instead of Ca^2+^, PedH showed a similar activity pattern as PedE (**Table 1**), but exhibited about 2-fold higher specific activities. Further, the functional concentration range of the cation cofactor differed dramatically for the two enzymes (**Fig. S2A**). While PedE showed enzyme activity at concentrations from 10 μM to 10 mM CaCl_2_ with a peak in activity at 1 mM, PedH activity was found with lanthanide concentrations as little as 10 nM and up to 100 μM with a peak in activity at 1 μM. From these data we calculated the dissociation constants *(K_D_)* for various metals to the corresponding enzyme and found that PedH has an 850- to 2500-fold higher binding affinity for lanthanides *(K_D_* = 25 – 75 nM; **Fig. S2B2**) as PedE does for Ca^2+^ *(K_D_* = 64 μM; **Fig. S2B1**). The subsequent determination of kinetic parameters with ethanol showed that *V_max_* of PedH was approximately 1.7-fold higher than that observed for PedE (10.6 U mg^−1^ *vs*. 6.1 U mg^−1^; **Fig. 3**). However, the corresponding *K_M_* was 2-fold lower for PedE compared to PedH (85 μM vs. 177 μM). A similar pattern was found with acetaldehyde and 2-phenylethanol, but with approximately 1.6-fold (2-phenylethanol) and 10–15-fold (acetaldehyde) lower catalytic efficiencies compared to those measured with ethanol. Statistical analysis (two-tailed t-test; *α* = 0. 05; *N* = 3; GraphPad Prism, version 7.03) revealed that all maximal velocities *(V_max_)* and binding constants *(K_M_)* except for the *K_M_* with ethanol were significantly different (*P* < 0.05) between PedE and PedH, however no significant differences could be observed in the catalytic efficiencies *(k_cat_/K_M_)*.

**FIG 3:**
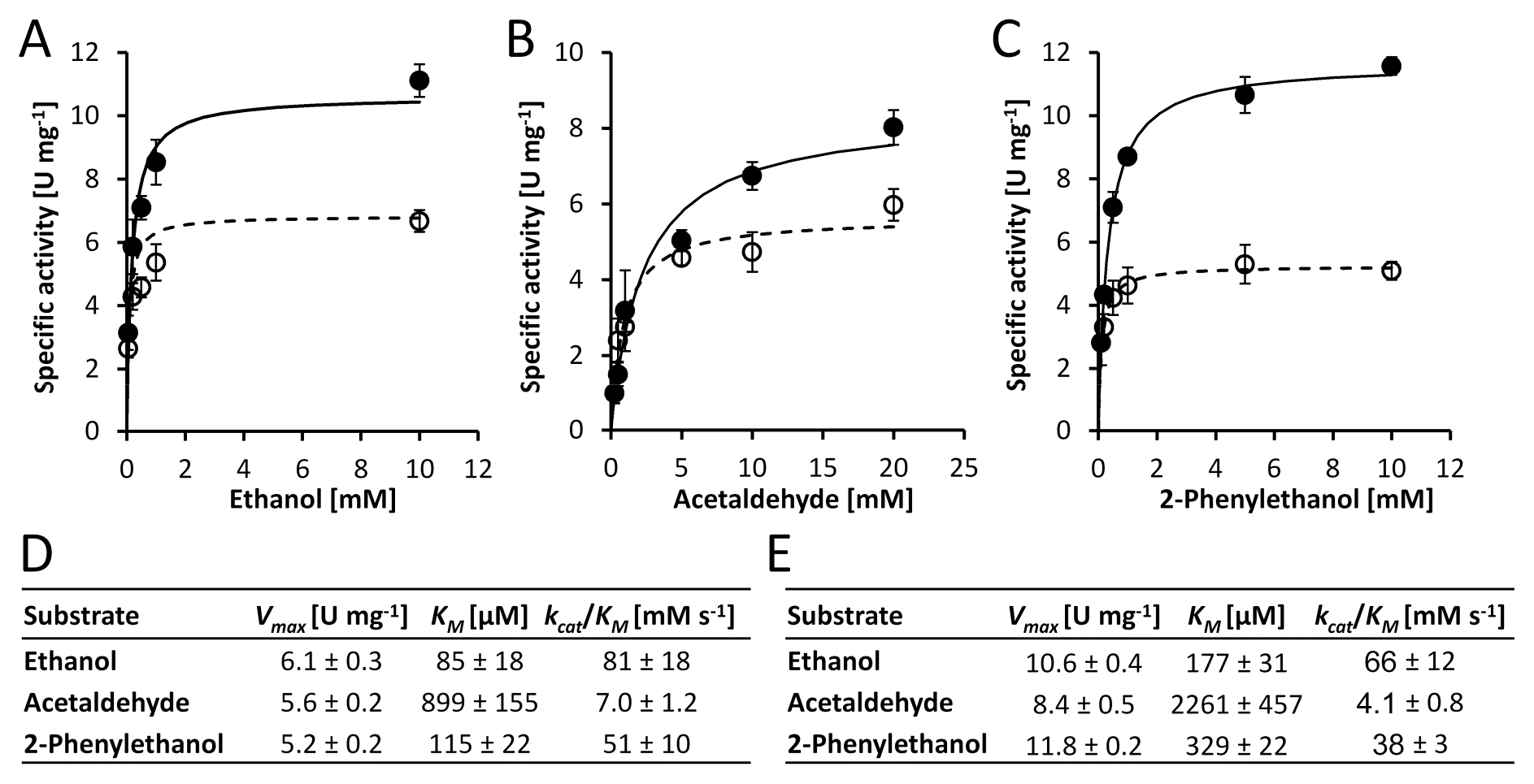
Kinetic parameter determination. **A-C**) Michaelis-Menten plot showing specific enzyme activities of PedE *(black circles)* and PedH *(white circles)* over a varying concentration of ethanol (**A**), acetaldehyde (**B**), and 2-phenylethanol (**C**). For PedE, 1 mM of CaCl_2_ and 50 μM PQQ was used, while for PedH, 1 μM PrCl_3_ and 1 μM PQQ was used in the reaction mixture. The data are given as mean value of triplicate measurements with error bars representing the standard deviation. Maximum velocity *(V_max_)*, substrate affinity (K_M_) and the catalytic efficiency *(k_cat_/K_M_*, were *k_cat_* is the turnover frequency per cofactor molecule) of PedE (D) and PedH (E) for different substrates are derived from **A-C** using nonlinear regression to a Michaelis-Menten model *(continuous lines* for PedH and *dashed lines* for PedE). Kinetik constants are represented as best fit values ± standard error.

### Growth with volatile alcohols in the presence and absence of lanthanides

In a next step, individual (*ΔpedE, ΔpedH, Δpqq*) and combinatorial (*ΔpedEΔpedH*) deletion mutants were tested for growth with several VOCs using an agar plate assay in the presence and absence of 20 μM lanthanum (**Fig. 4A**). Strains KT2440 (type strain), KT2440* (*Δupp* strain used as the parental strain for knockout mutants) and *ΔpedH* grew efficiently with ethanol, 1-butanol, and 2-phenylethanol in the absence of La^3+^. Strain *ΔpedE* displayed no growth under this condition. Even more interestingly, the addition of 20 μM of La^3+^ to the agar medium not only resulted in growth of the *ΔpedE* strain, but also restricted the growth of *ΔpedH*. The double mutant *ΔpedEΔpedH* and the *Δpqq* mutant, which is deficient in PQQ biosynthesis, showed no growth under both conditions. These experiments revealed that efficient growth with all tested alcohols, except the microbial fermentation product 2,3-butanediol, was dependent on the functional expression of PedE or PedH.

**FIG 4:**
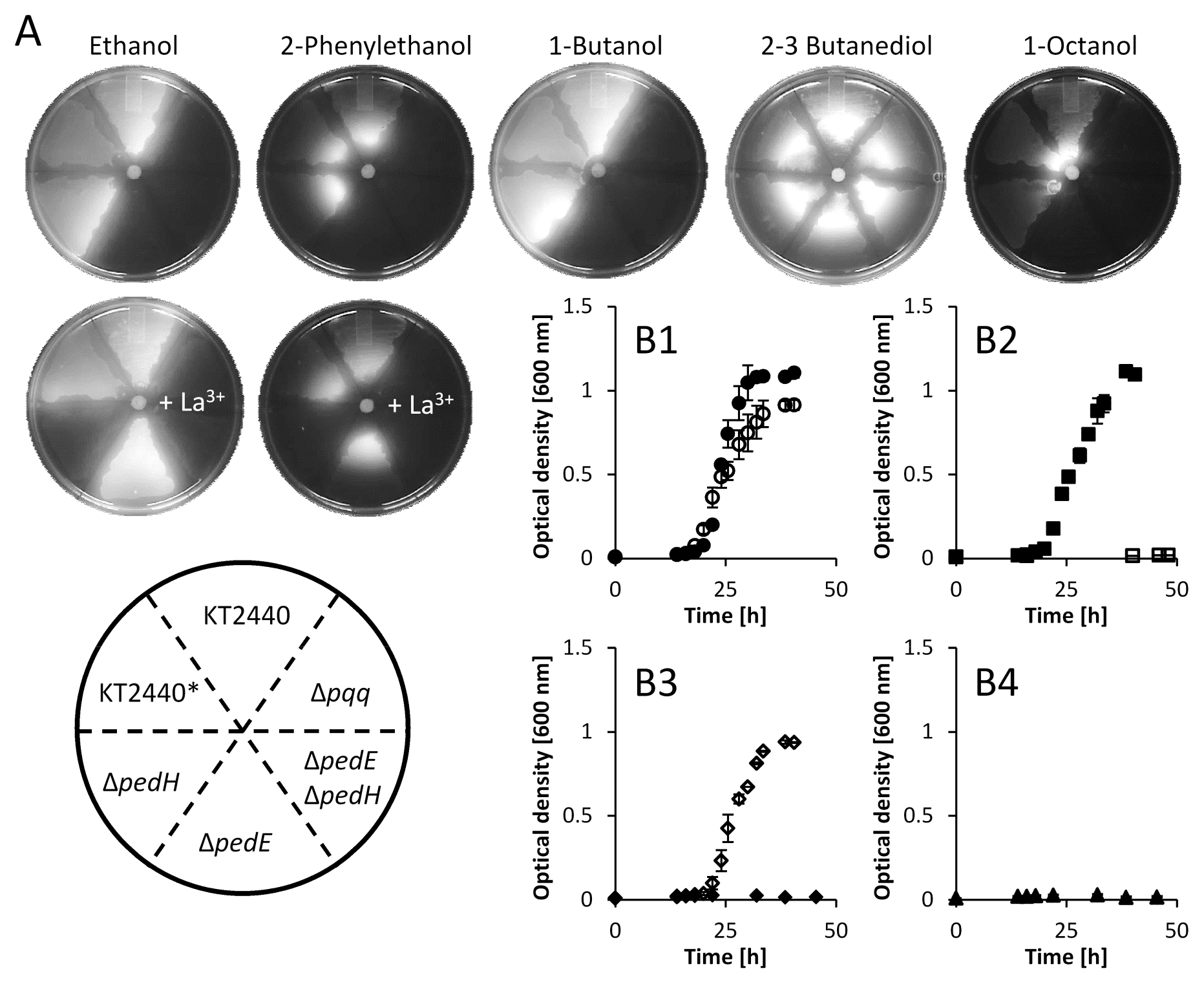
**A**) Growth with various substrates (10 μL drop of a 1:1 mix with DMSO) on M9 agar plates. Growth was quantified with a digital imaging system after 48 h using combined white light and UV illumination (ex. 254 nm). All pictures were sized, isolated from the background, and corrected for sharpness (+ 50%), brightness (+ 20%), and contrast (+ 40%). **B1–4**) Growth of KT2440* (Δupp strain used as the parental strain for knockout mutants; **B1**, *circles), ΔpedE* (**B2**, *diamonds), ΔpedH* (**B3**, *squares)*, and *ΔpedEΔpedH* (**B4**, *triangles)* in M9 medium with 5 mM 2-phenylethanol in the absence *(black symbols)* or presence of 20 μM La^3+^ *(open symbols)*. Growth was performed in 25 mL (125 mL plastic Erlenmeyer) at 30°C and 180 rpm shaking (Multifors) and quantified by optical density measurements at 600 nM. Data represent the mean of two individual cultures with error bars representing the corresponding range. All error bars are depicted but might not be visible due to the size of the corresponding symbol used for the mean value.

To validate these findings, growth experiments in liquid M9 medium with 2-phenylethanol as sole carbon and energy source in the presence and absence of 20 μM La^3+^ were performed (**Fig. 4B1–4**). For this, plastic Erlenmeyer flasks were used to avoid potential contaminations of rare earth metals (REM) from the glassware (33). Growth of the liquid cultures followed a similar pattern to that observed in the agar plate assay. While strain KT2440* (**Fig. 4B1**) showed growth after an 18–20 h lag phase and a peak in optical density at about 35 h in both conditions, the absence and the presence of lanthanum, strain *ΔpedEΔpedH* (**Fig. 4B4**) did not display growth in either condition. Growth of *ΔpedE* (**Fig. 4B3**) was observed exclusively in the presence of lanthanum. Lastly, strain *ΔpedH* (**Fig. 4B2**) showed growth similar to that of the KT2440* wildtype in the absence of lanthanum, but no growth was detected when 20 μM La^3+^ were supplemented.

### Transcriptional regulation of pedE and pedH determines growth with alcoholic volatiles

The previous experiments proved that for efficient growth with various VOCs the functional expression of at least one of the PQQ-ADHs is essential. The growth inhibition of the *ΔpedH* strain in the presence of lanthanum indicated a potential repression of the *pedE* gene in the presence of lanthanides, similar to recent reports in different methylotrophic bacteria (41–43). To proof this hypothesis, we constructed two reporter strains suitable for probing *pedE* and *pedH* promoter activities in KT2440*. When these strains were tested with 1 mM of 2-phenylethanol in M9 medium (**Fig. 5A**), the addition of up to 10 nM La^3+^ did not affect *pedE* promoter activity compared to the condition in the absence of lanthanum. In contrast, the presence of 100 nM – 100 μM of La^3+^ resulted in reduced *pedE* promoter activity. An inverse pattern was found for the *pedH* promoter. Here, very low activities were detected in the presence of up to 10 nM La^3+^. Upon addition of 100 nM or more lanthanum, expression from pedE promoter was induced with a peak at 10 μM.

**FIG 5:**
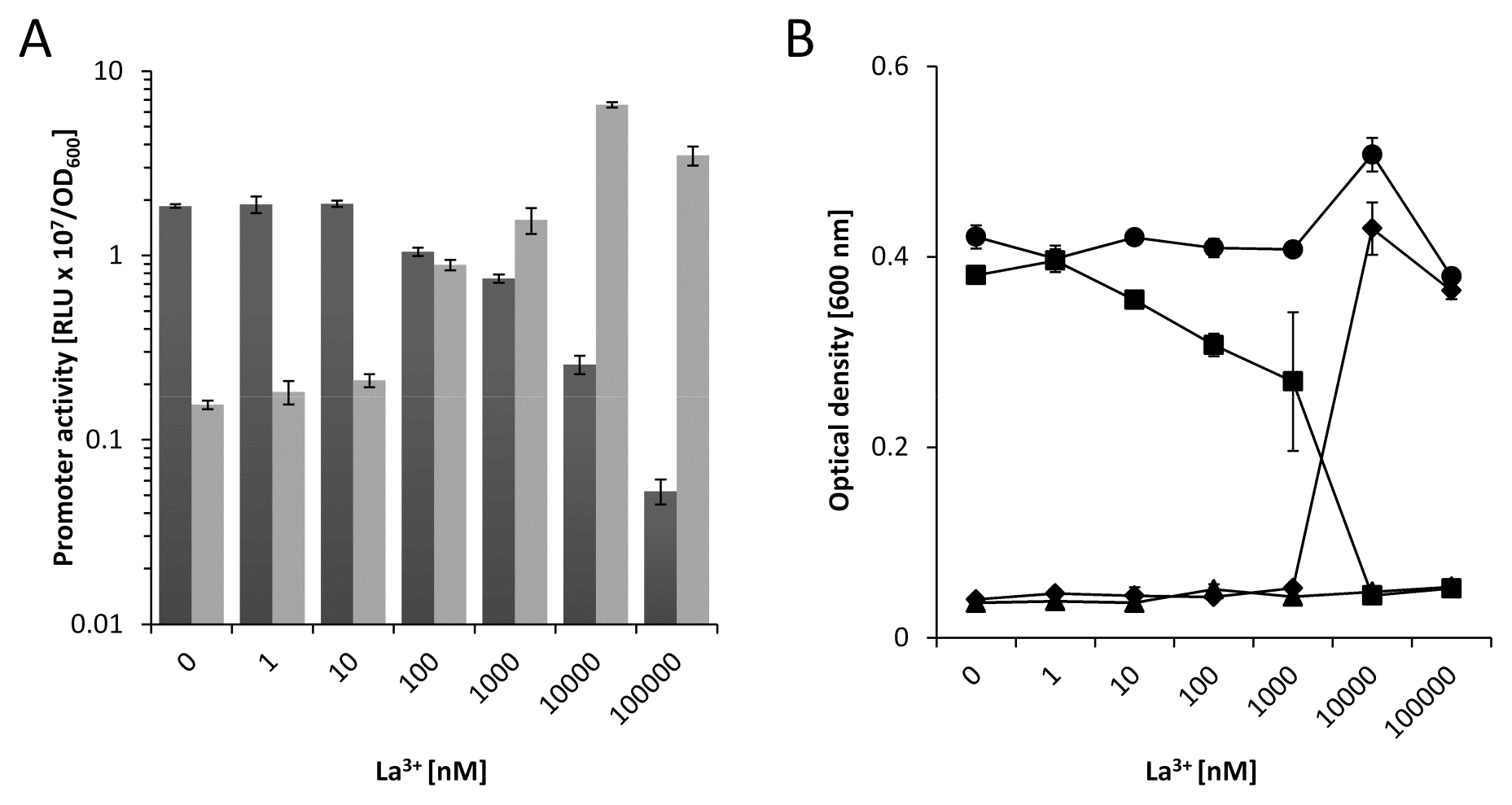
**A**) Activities of the *pedE (dark grey bars)* and *pedH (light grey bars)* promoters in strain KT2440* during incubation in liquid M9 medium (**A**) supplemented with 1 mM of 2-phenylethanol in the presence of varying concentrations of La^3+^. Promoter activities are presented as relative light units (RLU × 10^7^) normalized to OD_600_. **B**) Growth of KT2440* *(black circles), ΔuppΔpedE (black diamonds), ΔuppΔpedH (black squares)* and *ΔuppΔpedEΔpedH (black triangles)* in liquid M9 medium with 5 mM of 2-phenylethanol in the presence of different concentrations of La^3+^. Growth was determined as the optical density at 600 nm after incubation at 30°C for 24 h. Data are presented as mean values from biological triplicates and error bars represent the corresponding standard deviation.

The importance of transcriptional regulation of *pedE* and *pedH* was further tested by growth experiments with 2-phenylethanol in liquid M9 medium (**Fig. 5B**). Growth of KT2440* was not affected by the addition of up to 100 μM of lanthanum. On the other hand, strain *ΔpedH* showed a linear decrease in growth in the presence of increasing La^3+^ concentrations up to 1 μM, and no measurable growth when 10 μM or more La^3+^ was present in the medium. In contrast, growth of strain *ΔpedE* was only observed in the presence of 10 μM or more La^3+^. Strain *ΔpedEΔpedH* did not grow under any of the tested conditions. A similar correlation between growth and promoter activity of *pedE* and *pedH* was observed for Ce^3+^, Pr^3+^, Nd^3+^, and Sm^3+^ (**Fig. S3**).

These results demonstrate that KT2440 inversely regulates *pedE* and *pedH* promoter activity in response to varying lanthanide concentrations and suggests that this regulation represents the primary determinant for growth. However, in comparison to earlier studies with *M. extorquens* AM1, the effective lanthanide concentration needed for growth was much higher (10 μM *vs*. 5 nM)(41). As lanthanides are known to form very poorly soluble complexes with phosphate and hydroxide ions, we wondered whether this difference was caused by the minimal medium used for growth (MP vs. M9). When the experiments were repeated with MP medium, the same general trend and correlation of promoter activity and growth was found as described for M9 medium (**Fig. 6AB**). However, one difference was that the effective lanthanum concentration to trigger a transcriptional response and growth was considerably lower (1 nM and 10 nM). Another difference was that at a concentration of 10 nM La^3+^ minimal growth of both single mutant strains was observed. The latter observation indicated that environmental conditions might exist in which PedE and PedH are both functionally produced. To further test this hypothesis, an additional growth experiment was performed and indeed showed growth of both single mutants in a concentration range of 1 – 15 nM La^3+^ after prolonged (48 h) incubation (**Fig. S9**).

**FIG 6:**
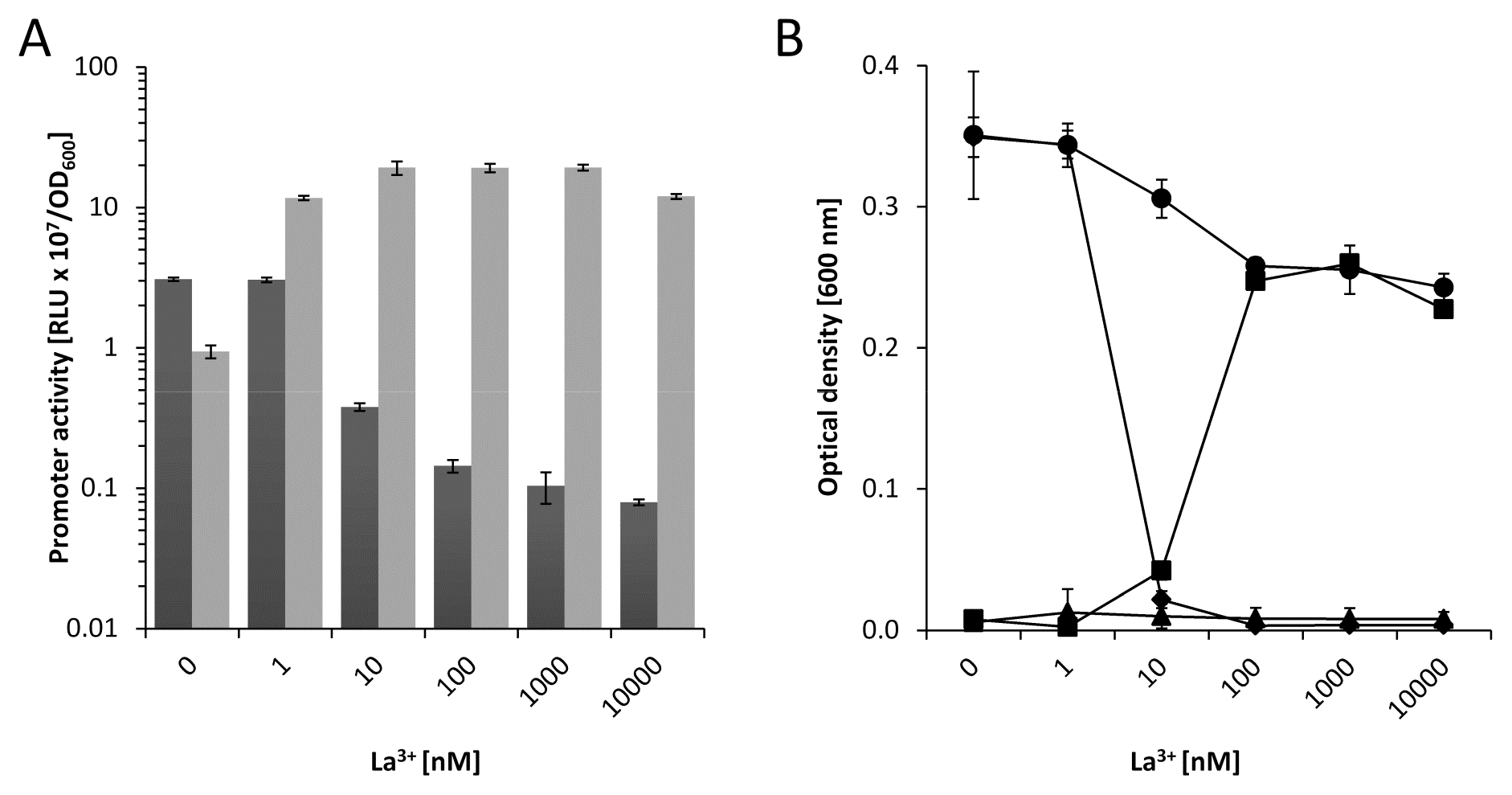
**A**) Activities of the *pedE (dark grey bars)* and *pedH (light grey bars)* promoters in strain KT2440* during incubation in liquid MP medium (**A**) supplemented with 1 mM of 2-phenylethanol in the presence of varying concentrations of La^3+^. Promoter activities are presented as relative light units (RLU × 10^7^) normalized to OD_600_. **B**) Growth of KT2440* *(black circles), ΔuppΔpedE (black diamonds), ΔuppΔpedH (black squares)* and *ΔuppΔpedEΔpedH (black triangles)* in liquid MP medium with 5 mM of 2-phenylethanol in the presence of different La^3+^ concentrations. Growth was determined as the optical density at 600 nm after incubation at 30°C for 24 h. Data are presented as the mean values from biological triplicates and error bars represent the corresponding standard deviation.

### Impact of PedE and PedH on transcriptional regulation

In *M. extorquens* AM1, the transcription of methanol dehydrogenases is regulated, at least partially, by the PQQ-dependent enzymes themselves (44). To test whether a similar outside-in signaling is also present in *P. putida* KT2440, expression from the *pedE* and *pedH* promoter were quantified during growth with 2-phenylethanol in MP medium (**Fig. 7**). In the absence of lanthanum, the *ΔpedH* strain showed a 4-fold induction of *pedE* promoter activity, whereas the *ΔpedE* strain exhibited a slight decrease (0.5-fold) in expression from the *pedE* promoter compared to KT2440* (**Fig. 7A**). The presence of 10 nM La^3+^ resulted in a strong reduction of *pedE* promoter activity in all strains (21-fold for KT2440*, 6-fold for *ΔpedE*, 127-fold for *ΔpedH)* compared to the control without lanthanum. In comparison to *pedE*, expression from the *pedH* promoter was considerably lower for all strains in the absence of lanthanum (**Fig. 7B**). However, when lanthanum was present, a strong induction of *pedH* promoter activity in strain KT2440* (37-fold) and *ΔpedE* (29-fold) was detected. Notably, the expression from the *pedH* promoter was dramatically reduced (2-fold vs. 37-fold) in the *ΔpedH* strain in comparison to the strains capable of producing a functional PedH protein.

**FIG 7:**
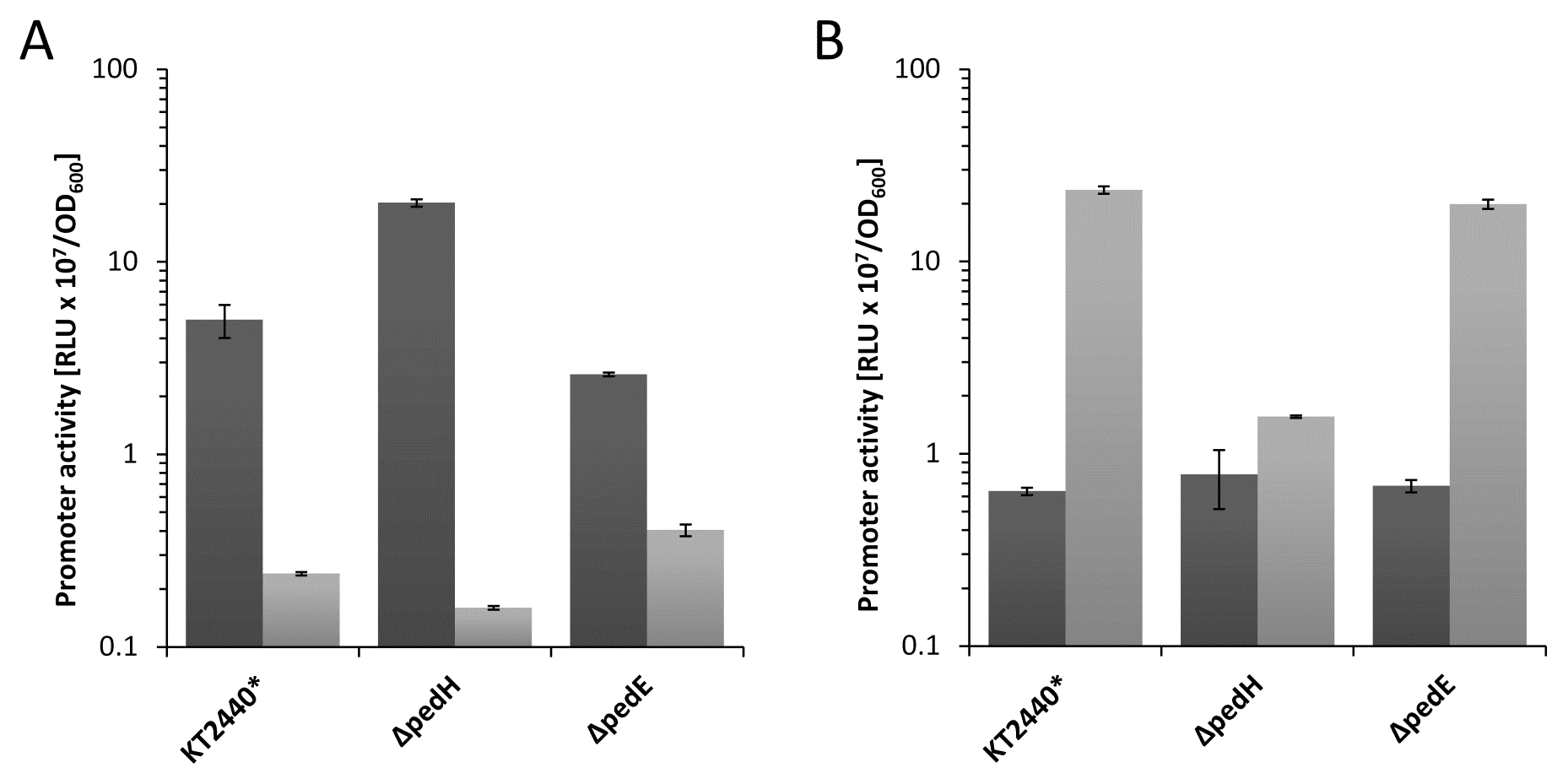
Activities of the *pedE* (**A**) and *pedH* (**B**) promoters in strains KT2440* (Δupp strain used as the parental strain for knockout mutants), *ΔpedE, ΔpedH and ΔpedEΔpedH* in liquid MP medium with 1 mM of 2-phenylethanol in the absence *(dark grey bars)* or presence of 10 nM La^3+^ *(light grey bars)*. Promoter activities are presented as relative light units (RLU × 10^7^) normalized to OD_600_. Data are presented as the mean values from biological triplicates and error bars represent the corresponding standard deviation. The promoter activities for each strain in the presence and absence of lanthanides were statistically analyzed (two-tailed *t*-test; *α* = 0.05; *N* = 3; GraphPad Prism, version 7.03) and found to be significantly different (*P* < 0.01).

## DISCUSSION

Lanthanide-dependent enzymes have so far been found exclusively within methylotrophic organisms (16, 28–30, 32, 33). Using purified enzymes, we uncover that PedH, one of the two PQQ-dependent ADHs (PQQ-ADHs) produced by the non-methylotrophic model organism *P. putida* KT2440, is also a lanthanide-dependent enzyme, which utilizes La^3+^, Ce^3+^, Pr^3+^, Nd^3+^, Sm^3+^, Gd^3+^ and Tb^3+^ as metal cofactor. The highest catalytic rates were observed with Pr^3+^ and Nd^3+^. Notably, with lanthanides of higher atomic masses than Nd^3+^ the specific activity decreased gradually, eventually resulting in no detectable activity with the heaviest lanthanides tested (Er^3+^ and Yb^3+^). An analogous effect was previously reported by Pol *et al.*, who investigated the impact of different lanthanides on growth rates of *Methylacidiphilum fumariolicum* SolV (33). A possible explanation for these observations is that the decreased atomic radius, which is a consequence of the lanthanide contraction, limits the more heavy lanthanides from being functionally incorporated into the active site of PedH (45).

Kinetic parameters determined with Pr^3+^ and the three model substrates ethanol, acetaldehyde and 2-phenylethanol revealed that *V_max_* of PedH is about twofold higher compared to its Ca^2+^-dependent counterpart PedE. It has been proposed that the increased activity can be explained by the higher Lewis acidity of the lanthanides in comparison to calcium (46). Interestingly, we found that the *K_M_* values of the lanthanide-dependent enzyme PedH with acetaldehyde or 2-phenylethanol are significantly smaller than those of the Ca^2+^-dependent protein PedE. One possible explanation for this result is the higher polarity that arises in the active pocket of the trivalent cation coordinating PedH compared to the divalent cation coordinating PedE. Another explanation could be a smaller catalytic pocket, due to the higher atomic radius of Pr^3+^ compared to Ca^2+^. The latter argument was proposed in an earlier study with the mxaF-type MDH of *M. extorquens* AM1 as a reason for decreased activities when Ca^2+^ was replaced with Ba^2+^ (47). In any case, apart from the approximately 2-fold increased specific activity of PedH compared to PedE, the substrate scope and catalytic efficiencies *(k_cat_/K_M_)* of both enzymes were found to be similar, which suggests that both enzymes are functionally redundant and only differ in their cofactor dependency.

Functional redundancy is a well-known mechanism to improve robustness in complex systems (48). The fact that many organisms express multiple PQQ-ADHs can be interpreted as an adaptation to maintain an important function under variable environmental conditions or in different microhabitats. Our study is supportive of such a hypothesis, as we demonstrate that under conditions of high lanthanide availability, efficient growth of cells with various naturally occurring alcoholic VOCs relies on the functional production of the lanthanide-dependent ADH PedH. Similarly, for growth in the absence of lanthanides, functional production of the calcium-dependent ADH PedE is mandatory. In this context it is important to note that growth in the agar plate assay used in this study is restrictive, as it depends on diffusion and evaporation of the volatile substrates. Thus, the assay most likely cannot discriminate between substrates for which PedE or PedH function are essential and substrates for which other but less efficient catabolic routes exist. We found that growth on ethanol (unpublished data) and 1-butanol (19), but not 2-phenylethanol (current study) is possible, but less efficient for a strain lacking PedE and PedH. From our and published data from a previous study (49), we therefore conclude that beside the fact that PedE and PedH is not essential for growth with short chain aliphatic alcohols (C2 to C5), both enzymes provide rapid conversion of these substrates, which is crucial for efficient growth under restrictive conditions.

We further show that growth phenotypes strongly correlate with inverse transcriptional regulation of *pedE* and *pedH*. Similar results have been reported for several methylotrophic bacteria (41, 42). When cells of *P. putida* KT2440 were grown in liquid MP medium, the addition of as little as 10 nM of lanthanides was sufficient to trigger *pedE* repression and a strong (20-fold) concomitant induction of *pedH*. The transcription of *pedH* was found to be strongly influenced by the PedH protein itself, implying a role for PedH as a lanthanide sensory module. In *M. extorquens* AM1 the transcription of the calcium-dependent methanol dehydrogenase *mxaF* strictly relies on the presence of XoxF proteins (41, 44). Our data demonstrate that this regulation is different in *P. putida*, as *pedE* is only partially repressed by PedH. The fact that *pedE* repression in the presence of lanthanum is still observed in the *ΔpedH* mutant strain, together with the notion that the induction of *pedH* is not fully mediated by PedH, strongly suggests the existence of at least one additional lanthanum-responsive regulatory module. Notably, a very recent study identified that the transmembrane associated sensory histidine kinase MxaY mediates the lanthanide-responsive switch of the PQQ-dependent MDHs in *Methylomicrobium buryatense* (43). In *P. putida* KT2440 three different membrane associated sensory histidine kinases, PedS1 (PP_2664), PedS2 (PP_2671) and PP_2683, are encoded within close proximity of *pedE* and *pedH* as part of the predicted ErbR (AgmR) regulon (17). Whether one of these sensor kinases serves a similar function as MxaY needs to be determined in future studies.

From an ecological point of view, it is interesting that growth and inverse regulation occur even in presence of high Ca^2+^ concentrations (100 μM) when only nanomoles of lanthanum are supplied. From these data we conclude, similar to previous studies with *M. buryatense* or *M. extorquens* (32, 41), that the lanthanide-dependent enzyme PedH is the preferred PQQ-ADH when both metal cofactors are simultaneously accessible. Nevertheless, it was also demonstrated that under certain low REM concentrations, specific conditions exist in which both single mutants can grow. This suggests that the inverse regulation of the two enzymes is not a strict *on-off* switch, but rather operates by strongly shifting the transcription in favor of one of the enzymes depending on the REM concentration.

REM utilizing PQQ-ADHs have been suggested to be ancestral and more widespread than their calcium-dependent homologues (15, 50). This might indicate that calcium-dependent enzymes have evolved to colonize different and/or additional environmental niches in which lanthanide availability is less pronounced. Compared to soil environments, especially the rhizosphere, lanthanide concentrations in the phyllosphere and endosphere, as well as in other non-plant higher organisms are comparably low (51–54). It is thus tempting to speculate, that Ca^2+^-dependent enzymes are of particular relevance for interactions with multicellular organisms outside of soil environments.

Metabolic interdependencies have been proposed as driving force for species co-occurrence and the emergence of mutualism in diverse microbial communities impacting their robustness, structure, and function (55–57). This is of particular interest in the context of periplasmic PQQ-ADHs, as organic alcohols and the corresponding oxidation products are not only crucial intermediates of the global carbon cycle, but can also exhibit additional functions including signaling and growth inhibition (4–7, 58). A recent study reported that regulation of the MxaF- and XoxF-type MDHs in a methanotrophic bacterium can be influenced by the presence of a non-methanotrophic methylotroph in co-culture experiments (59). The authors nicely demonstrate that during co-cultivation in the presence of methane and lanthanides, the methanotrophic bacterium shifts its gene expression from the *xoxF*- to the *mxaF*-type MDH. As a result of this change, leakage of methanol from the methanotroph was observed, which subsequently served as growth substrate for the non-methanotrophic partner. Although the mechanism of this phenomenon is not yet resolved, it indicates that different types of PQQ-ADHs might not only be important for potential interactions with higher organisms as discussed above, but also within microbial communities. Based on our data, one can speculate that similar interactions are not limited to methano- and methylotrophic bacteria, but are relevant in a much broader ecological context.

Lastly, the discovery of lanthanides as a cofactor in biotechnological important organisms other than methylotrophic bacteria expands the possible applications one can envision for biomining, bioleaching, and recycling processes of rare earth metals (60–65). As such, we believe that future research about lanthanide-utilizing enzymes and organisms will improve our understanding of natural and synthetic microbial communities and could provide a basis for novel biotechnological tools and processes.

## MATERIAL AND METHODS

### Bacterial strains, plasmids, and culture conditions

Strains and plasmids used in this study (**Table S1**) as well as a detailed description of their construction (**Text S1**) can be found as supplementary material. Unless otherwise noted, *Pseudomonas putida* KT2440 and *Escherichia coli* strains were maintained on solidified (1.5% agar [w/v]) Lysogeny Broth (LB, Maniatis *et al.*, 1982). Strains were routinely cultured in liquid LB medium, a modified M9 salt medium containing 74 mM phosphate buffer (pH 7), 18.6 mM NH_4_Cl, 8.6 mM NaCl, 2 mM MgSO_4_, 100 μM CaCl_2_ with a trace element solution containing Na_3_-citrate 51 μM, ZnSO_4_ 7 μM, MnCl_2_ 5 μM, CuSO_4_ 4 μM, FeSO_4_ 36 μM, H_3_BO_3_ 5 μM, NaMoO_4_ 137 nM, NiCl_2_ 84 nM or a modified MP medium (67) containing 100 μM CaCl_2_ instead of 20 μM CaCl_2_ supplemented with succinate or 2-phenylethanol as sole source of carbon and energy at 30°C with shaking. Where indicated, 40 μg mL^−1^ kanamycin or 15 μg mL^−1^ gentamycin for *E. coli* and 40 μg mL^−1^ kanamycin, 20 μg mL^−1^ 5-fluoro uracil or 30 μg mL^−1^ gentamycin for *P. putida* strains was added to the medium for maintenance and selection, respectively.

### Growth experiments in liquid medium

All liquid growth experiments were carried out using a modified M9 minimal salt medium or MP medium (see above) supplemented with 25 mM succinate or 5 mM 2-phenylethanol as carbon and energy source. To avoid potential lanthanide contaminations from glassware, all growth experiments were carried out in 125 mL polycarbonate vessels (Corning) or in polypropylene 96 well 2 ml deep well plates (Carl Roth). If not stated otherwise, precultures were grown in 5 ml minimal medium (15 mL Falcon tubes) supplemented with succinate at 30 °C and 180 rpm using a rotary shaker (HT Minitron, Infors). The next day, cultures were washed three times in fresh minimal medium without a carbon and energy source and used to inoculate 1 ml (for 2 ml deep well plates) or 25 ml (for 125 ml polycarbonate vessels) of fresh medium to an initial optical density at 600 nm (OD_600_) of 0.01. Subsequently, cultures were supplemented with the carbon and energy source as well as varying concentrations of lanthanides and incubated at 30°C and 180 rpm (for 125 ml polycarbonate vessels) or 800 rpm (for 2 ml deep well plates). For experiments in 125 ml polycarbonate vessels, growth was monitored by measuring the OD_600_ at regular intervals using a photometer (BioPhotometer, Eppendorf). For experiments carried out in 2 ml deep well plates, OD_600_ was determined after 24 h or 48 h by measuring 200 μL of cell culture transferred to a microtiter plate (Greiner bio-one) in a microplate reader (POLARstar Omega, BMG Labtech). All data are presented as the mean value of biological triplicates with error bars representing the corresponding standard deviation.

### Agar plates assay

For growth on solidified medium (1.5% agar [w/v]) with different substrates, M9 medium plates without addition of a carbon source and trace element solution were freshly prepared with or without the addition of 20 μM lanthanum chloride. Cell mass of the strains was obtained from LB agar plates, suspended in M9 medium without carbon and energy source, and adjusted to an optical density of 0.5. After drying the plates for 20 minutes in a laminar flow cabinet, 10 μL of each cell suspension was dropped onto the same plate and distributed using an inoculation loop on about 1/6 of the plate’s surface. When all strains were distributed, a 10 μL drop of a 1:1 mixture (v/v) of ethanol, 1-butanol, 2–3 butanediol, 1-octanol, or 2-phenylethanol in dimethyl sulfoxide (DMSO) was placed in the middle of the plate. Subsequently, the plates were sealed in plastic bags and incubated at room temperature. After 48 h, growth was quantified with a digital imaging system (Vilber Lourmat, QUANTUM ST4) using the standard fluorescence settings with combined white light and UV illumination (ex. 254 nm) for 1 sec, an aperture of 11 and using the preinstalled F590 nm filter. All individual pictures were subsequently sized, isolated from the background, and corrected for sharpness (+ 50%), brightness (+ 20%), and contrast (+ 40%) using the graphic formatting function in Microsoft PowerPoint.

### Transcriptional reporter assays

For transcriptional reporter assays, *P. putida* harboring either Tn7-based *pedE-lux* and *pedH-lux* transcriptional reporter fusion were grown overnight in a modified MP medium with succinate, washed three times in MP medium with no added carbon source and finally suspended in MP medium or M9 medium with 1 mM 2-phenylethanol to an OD_600_ of 0.1. For luminescence measurements, 180 μl of cell suspension were added to 20 μl of tenfold concentrated metal salt solution in white 96-well plates with clear bottom (μClear, Greiner Bio-One). Microtiter plates were placed in a humid box to prevent evaporation, incubated at 30°C with continuous agitation (200 rpm) and light emission as well as OD_600_ were recorded at regular intervals in a FLX-Xenius plate reader (SAFAS, Monaco) for up to 6 h. For both parameters, background provided by the MP medium was subtracted, and the luminescence was normalized to the corresponding OD_600_. Experiments were performed in biological triplicates and data are presented as the mean value with error bars representing the corresponding standard deviation.

### Enzymatic assays

Details about the expression and purification procedure for PedE and PedH can be found as supplementary material (**Text S1**). Enzyme activities of purified PedE and PedH were measured using a dye-linked colorimetric assay in 96 well microtiter plates (Greiner bio-one) based on previous studies (24, 68). Under optimized conditions (**Fig. S4-S8**), one well contained a total volume of 250 μL of assay solution supplemented with: 100 mM Tris HCl pH 8; 500 μM PMS; 150 μM 2,6-dichlorophenol indophenol (DCPIP); 25 mM imidazole; 1 mM CaCl_2_ for PedE or 1 μM PrCl_3_ for PedH; 1 μM PQQ for PedE or 50 μM PQQ for PedH; 12.5 μL substrate and 2.5–20 μg/ml enzyme. The reaction was started by addition of the substrate to the reaction mixture and the activity was calculated based on the change of OD_600_ within the first minute upon substrate addition. The molar extinction coefficient of DCPIP was experimentally determined to be 24.1 cm^−1^M^−1^ at pH 8 (**Fig. S2**). Due to a substrate independent background activity, the assay solution without substrate was incubated for 45 minutes at 30°C prior to enzyme activity measurements. As activities were between 8- and 12-fold higher when using imidazole compared to ammonium chloride or ethylamine, imidazole was used in all experiments (Fig. S3). Negative control reactions, including the potential effect of BSA or assay mixture without the addition of enzyme, did not show any reduction of DCPIP under the conditions used (data not shown). All assays were performed in three replicates and data are presented as the mean value with error bars representing the corresponding standard deviation.

### Metal dependency of the enzymes

To test the metal dependency of PedE and PedH, a similar set-up as described above was used omitting CaCl_2_ for PedE or PrCl_3_ for PedH in the assay solution. 1 μM of different rare earth metals was added prior to incubation at 30°C. These included LaCl_3_, CeCl_3_, PrCl_3_, NdCl_3_, SmCl_3_, GdCl_3_, ErCl_3_, YbCl_3_, ScCl_3_ and YCl_3_. Activities were determined in triplicates as described above.

### Enzyme kinetics

The kinetic constants of the enzyme substrate combinations were determined using the enzyme assay described above with various substrate concentrations measured in triplicates. The resulting activity constants were calculated by fitting the enzyme activities by nonlinear regression to the Michaelis-Menten equation using the ‘Michaelis-Menten’ least-square fit method with no constrains in GraphPad Prism (GraphPad Software, version 7.03).

### Homology models

The PedE and PedH homology models were built with Swiss-Model (69). As ExaA has the highest sequence similarity with both PedE (60%) and PedH (49%) of all available crystal structures in the Swiss-Model template library, the crystal structure of the PQQ dependent ADH ExaA of *P. aeruginosa* (1FLG) was used as a template for model construction (23). Visualization of the models was carried out using PyMOL (70).

## ACKNOWLEDGEMENTS

The authors would like to thank Lena Stehle and Svenja Moors for their help in the establishment of the enzymatic assay. Prof. Altenbuchner, Dr. Nadja Graf, Dr. Joanna Goldberg and Prof. Herbert Schweizer are acknowledged for providing different strains and plasmids. We further would like to thank Prof. Thorsten Thomas and Dr. Brendan Colley for critical reading of the manuscript draft and Prof. Bernhard Hauer for his continuous support. The authors further declare that the research was conducted in the absence of any commercial or financial relationships that could be construed as a potential conflict of interest.

